# A nuclear function for an oncogenic microRNA as a modulator of snRNA and splicing

**DOI:** 10.1101/2021.09.30.462508

**Authors:** Rachid EI Fatimy, Yanhong Zhang, Evgeny Deforzh, Mahalakshmi Ramadas, Harini Saravanan, Zhiyun Wei, Rosalia Rabinovsky, Nadiya M. Teplyuk, Erik J. Uhlmann, Anna M. Krichevsky

**Affiliations:** Department of Neurology, Brigham and Women’s Hospital and Harvard Medical School, Boston, MA 02115, USA.; Institute of Biological Sciences (ISSB-P), Mohammed VI Polytechnic University (UM6P), Benguerir 43150, Morocco.; Shanghai Key Laboratory of Maternal Fetal Medicine, Shanghai First Maternity and Infant Hospital, School of Medicine, Tongji University, Shanghai 200092, China

**Keywords:** miR-10b, glioblastoma, nucleus, U6 snRNA, splicing machinery, CDC42

## Abstract

miRNAs are regulatory transcripts established as repressors of mRNA stability and translation. Here we demonstrate that a key onco-miRNA, miR-10b, binds to U6 snRNA, a core component of the spliceosomal machinery. We provide evidence of direct binding between miR-10b and U6, *in situ* visualizations of miR-10b and U6 co-localization in glioma cells and patient-derived tumor tissues, and biochemical co-isolation of miR-10b with the components of the spliceosome. We further demonstrate that miR-10b modulates U6 N-6-adenosine methylation and pseudouridylation, U6 binding to splicing factors SART3 and PRPF8, and regulates U6 stability, conformation, and levels. These effects on U6 result in splicing alterations, illustrated by the altered ratio of the isoforms of a small GTPase CDC42, reduced overall CDC42 levels, and downstream CDC42 -mediated effects on cell viability. We, therefore, present an unexpected intersection of the miRNA and splicing machineries and a new nuclear function for a major cancer-associated miRNA.

## INTRODUCTION

MiR-10b is one of the miRNAs most strongly associated with various types of cancers (Lund, 2010, Tehler, Hoyland-Kroghsbo et al., 2011). Since the initial report of the aberrant miR-10b expression and association with breast cancer metastasis in 2007 (Ma, Teruya-Feldstein et al., 2007), multiple studies have linked miR-10b to cancer progression, regulation of specific cancer-promoting signaling pathways, and clinical outcomes in a broad spectrum of malignancies (Sheedy & Medarova, 2018). miR-10b has been since implicated in cancer cell proliferation, epithelial-mesenchymal transition, invasion, metastasis, angiogenesis, and drug resistance. In addition, an investigation of miR-10b-deficient mice demonstrated that this miRNA is essential in tumorigenesis (Kim, Siverly et al., 2016). These data led to active exploration of miR-10b as a therapeutic target as well as a cancer biomarker by academia and the biotechnology industry.

Perhaps the most distinct and intriguing is the activity of miR-10b in malignant gliomas, primary brain tumors affecting both adult and pediatric populations. While in most other cancers miR-10b primarily drives metastasis, in gliomas it emerged as a critical survival factor required for tumor cell viability (Gabriely, Yi et al., 2011b). The major difference in miR-10b activity in cancer cells of different lineages likely stems from its expression patterns: whereas expressed in normal extracranial tissues, miR-10b gene is completely silenced in the fetal and adult brain cortex but derepressed in the low- and high-grade gliomas (Gabriely, Yi et al., 2011b, Teplyuk, Uhlmann et al., 2016). This aberrant out-of-place expression and activity of miR-10b in the brain tumors become addictive for the tumor cells and leads to miR-10b dependence (El Fatimy, Subramanian et al., 2017). miR-10b inhibition is, therefore, considered a highly promising therapeutic approach for malignant gliomas, the uniformly lethal tumors presenting a major unmet need in current oncology.

Intriguingly, despite a decade of research efforts, key mRNA targets of miR-10b regulation remain largely controversial and often tumor- or even tumor subtype-specific (Gabriely, Teplyuk et al., 2011a, Sheedy & Medarova, 2018, Teplyuk, Uhlmann et al., 2015). Glioma studies indicate that direct miR-10b targets are poorly predicted by various bioinformatics algorithms and that miR-10b regulation is unconventional (Guessous, Alvarado-Velez et al., 2013, Teplyuk et al., 2016). Specifically, the miRNA “seed” (nucleotides 2-7 at the 5’ end of a miRNA), known as the primary determinant of miRNA binding and targeting, does not appear essential for the miR-10b activity. Neither *in vitro* experiments on glioma and glioma-initiating cells nor TCGA analyses support the computational predictions of miR-10b targets, challenging the mechanistic studies. Our prior work demonstrates that miR-10b regulates mRNA splicing, partly by controlling the levels of several splicing factors, including MBNL1-3 (Teplyuk et al., 2016).

Our observations of glioma-initiating cells, called glioma stem cells (GSCs), along with recent reports establishing the presence of Dicer, the major enzymatic complex involved in the final steps of miRNA maturation, in the GSC nuclei (Bronisz, Rooj et al., 2020), suggested additional, previously unexplored nuclear functions for miRNAs. The evidence of the non-conventional miRNA regulation, including nuclear and non-seed-mediated mechanisms, has been supported by recent studies (Zhang, Artiles et al., 2015). However, so far, it has attracted little attention in the studies of human cancer. To unravel a fundamental mechanism of glioma addiction to miR-10b, we performed an unbiased, RNome-wide analysis of the miR-10b interactome, not confined by the bioinformatic predictions and mapping to mRNAs. This strategy, based on the Covalent Ligation of Endogenous Argonaute-bound RNAs and their high-throughput RNA sequencing (CLEAR-CLIP) (Moore, Scheel et al., 2015), led to the discovery of U6 small nuclear RNA (snRNA), the core component of a spliceosome as the major miR-10b binding transcript. We further developed a range of complementary biochemical and imaging-based techniques to investigate a new and unanticipated miRNA function in controlling U6 snRNA metabolism, spliceosome assembly, and alternative splicing in the cancer cells. Therefore, this study further expands the network of miRNA regulation beyond mRNA and lncRNA targets and demonstrates the interface of miRNA and splicing machineries.

## RESULTS

### MiR-10b binds to snRNA U6 in glioma cells

To investigate endogenous miR-10b targets, we employed the modified CLEAR-CLIP protocol in glioma cells (Moore et al., 2015). The technique relies on miRNA crosslinking with proximal transcripts, ligation, and sequencing of chimeras. Therefore, it provides an unbiased mapping of the miRNA-target interactions, independent of bioinformatic predictions (Fig 1A). The identified miR-10b chimeras contained fragments matching multiple predicted and validated mRNA targets (Fig 1B, Dataset EV1). Of the identified miR-10b-binding transcripts, 115 corresponded to protein-coding and 31 to non-protein coding genes, and the majority did not contain miR-10b seed, despite the extensive base-pairing with miR-10b (Fig 1C and EV1, Dataset EV2). Surprisingly, the most common chimeras were formed by miR-10b and the sequences corresponding to U6 snRNA. Inspection of U6 snRNA transcript demonstrated extensive base pairing between its 3’ end and miR-10b, with a stretch of 10 complementary nucleotides at the positions 14-23 of miR-10b, and additional complementarity beyond this site (Fig 1D). To validate the binding between miR-10b and U6, we further used CLEAR-CLIP to produce miRNA chimeras from several miR-10b-expressing cancer cell lines (Fig EV2) and detect the product with a pair of PCR primers, one corresponding to miR-10b and another to U6. We detected miR-10b-U6 chimeras in glioma cell lines and GSCs, with trace levels also present in other cancerous non-glioma cells (Fig 1E). Collectively, these data demonstrate that miR-10b binds to U6 snRNA in various cancer cells, and this intermolecular interaction is prevalent in glioma cells.

**Figure 1.**
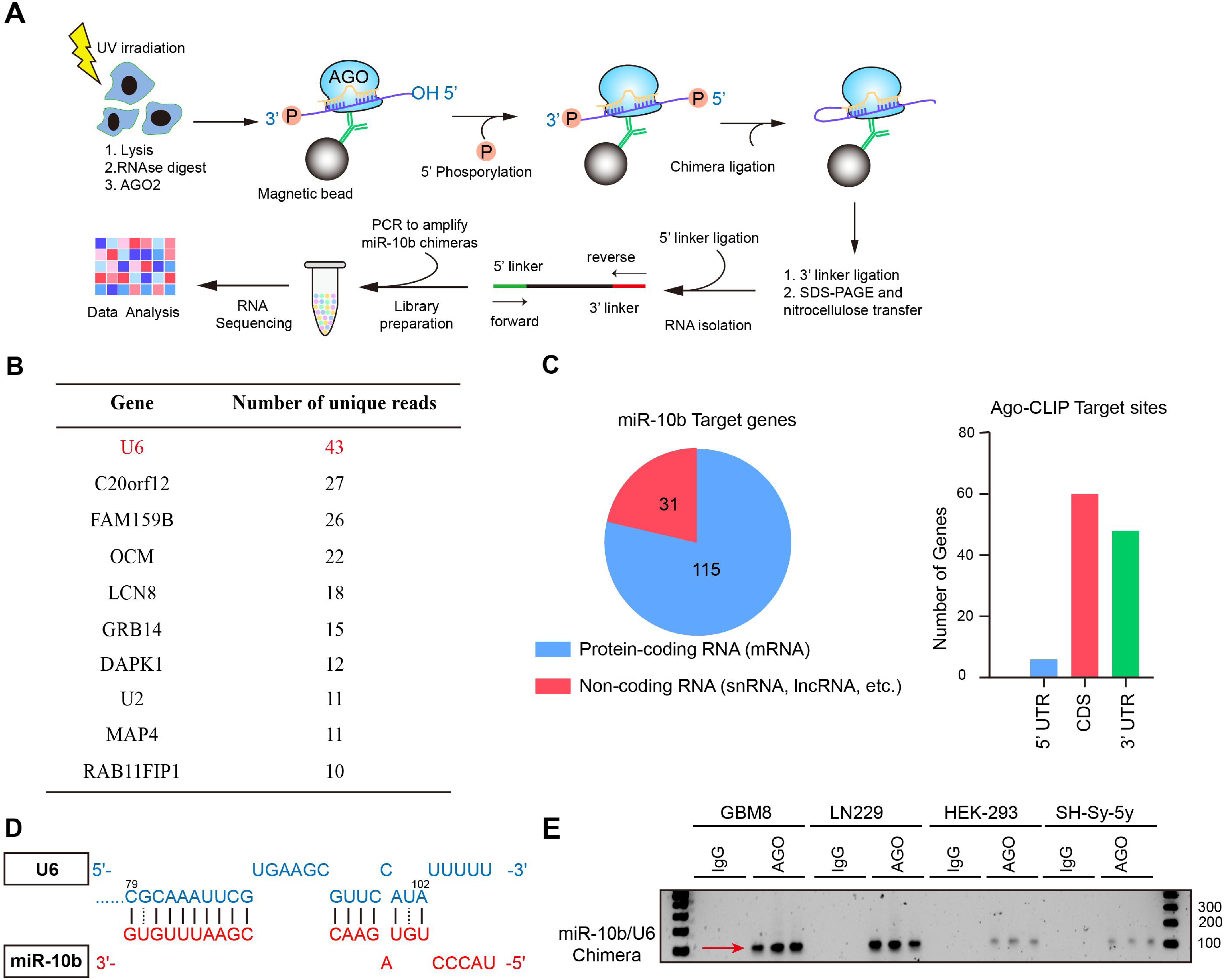
CLEAR-CLIP identifies U6 snRNA as the top miR-10b interacting transcript. A. Schematics of CLEAR-CLIP procedure, described in details in the Methods. B. Top genes represented by the highest number of unique reads identified in chimeric miR-10b libraries in LN229 cells. C. Summary of miR-10b chimeras and binding site distribution in mRNAs. More details are provided in Figure S1, Table S1 and Table S2. D. miR-10b binding site within U6 snRNA, at the 3’ end (positions 79 to 102) of U6. E. miR-10b-U6 chimera detected by PCR, with one primer corresponding to miR-10b and another to U6, in CLEAR-CLIP libraries produced from glioma LN229 cells, GBM8 stem cells, and non-glioma HEK-293 and SH-Sy-5y cell lines (n=3).

snRNA U6 is a major RNA component of the spliceosome. It localizes predominantly, if not exclusively, to the nuclear space (Spiller, Boon et al., 2007). Mature miRNAs are thought to operate mostly in the cytosolic compartment; however, in specific conditions some of them could be found in the nucleus (Hwang, Wentzel et al., 2007, Liu, Lei et al., 2018, Politz, Hogan et al., 2009). In agreement with this, in glioma cells, the major enzymatic complex responsible for the final step of miRNA processing, Dicer, is aberrantly localized to the nucleus (Bronisz et al., 2020). To investigate miR-10b subcellular localization, we fractionated glioma cells into cytosolic and nuclear fractions. More than half of the cellular miR-10b pool was confined to the nuclei; of this nuclear pool, >40% was associated with the insoluble fractions containing the spliceosomal complexes (Fig 2A).

**Figure 2.**
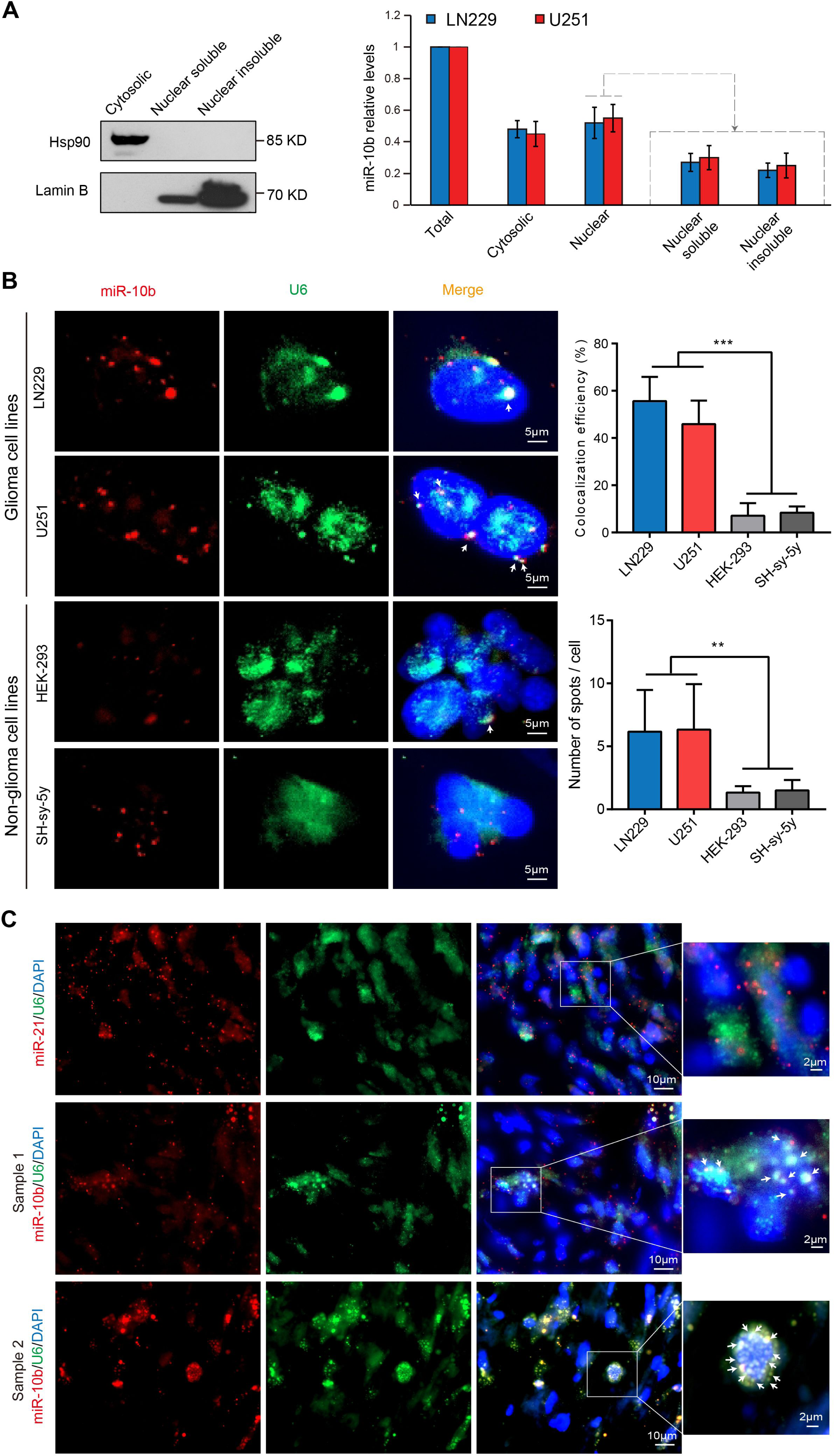
Co-localization of miR-10b and U6 snRNA in human GBM cells and tumors. A. Fractionation of glioma cells indicates that miR-10b is distributed in both cytosolic and nuclear compartments. Representative Western blotting of cytoplasmic, nuclear soluble, and nuclear insoluble fractions of glioma LN229 cells for Hsp90 (cytosolic) and Lamin B (nuclear) markers (left). MiR-10b levels in cytoplasmic, nuclear soluble, and nuclear insoluble fractions in LN229 and U251 glioma lines by qRT-PCR (mean ± SD, n=3) (right). B. Representative FISH images of miR-10b (Alexa FluorTM 546, red) and U6 (Alexa FluorTM 488, green) in cultured LN229 and U251 glioma cells and non-glioma HEK-293 and SH-Sy-5y cell lines with a fluorescently labeled probe, and nuclei stained with DAPI. Arrows mark the colocalization of miR-10b with U6. Quantification of the colocalization between miR-10b and U6 is presented in right panels. The percentage of cells with the colocalized miR-10b and U6 signals and the number of colocalization spots per cell are shown. At least 80 cells were analyzed for each cell line. *P* values were calculated using one-way ANOVA. ** *P* < 0.01; *** *P* < 0.001. C. Representative FISH images of miR-10b or miR-21 (red) and U6 (green) in patient-derived GBM tissues with the corresponding fluorescently labeled probes, and nuclei stained with DAPI. Arrows mark the colocalization of miR-10b with U6.

To further explore the association between miR-10b and snRNA U6 in cancer cells, we visualized mature miR-10b and U6 in LN229 and U251 glioma cells and other cancer cells using RNA *in situ* hybridization (FISH). miR-10b localized in multiple nuclear and perinuclear speckles and often co-localized with U6 snRNA (Fig 2B). The co-localization was more common in glioma than in other cell lines (Fig 2B). Importantly, patient-derived GBM tumors also exhibited remarkable, in some cells almost perfect colocalization between miR-10b and U6 (Fig 2C). No colocalization was detected between U6 and another highly abundant cancer-associated miRNA, miR-21, indicating the selectivity and sequence-specificity of the U6 binding to miR-10b.

### miR-10b is a part of the spliceosomal SART3 and PRPF8 RNPs

To further validate the binding between miR-10b and U6, we complemented the pool-down of Ago complexes with the similar RNA iCLIP technique employing various U6-interacting spliceosomal proteins as tags. Both U6 and miR-10b have been significantly enriched in the spliceosomal SART3 and PRPF8 complexes (Fig 3A and EV3A, B). Furthermore, miR-10b-U6 chimeras have been detected in the cross-linked and ligated SART3 and PRPF8 complexes, similar to the Ago complexes (Fig 3B) indicative of the miR-10b interaction with U6 snRNP in the splicing complex. SART3 or the spliceosome-associated factor 3 (aka hPrp24) is a core spliceosomal factor and a single spliceosomal protein that binds U6 selectively (Martin-Tumasz, Richie et al., 2011). It participates in the U6 biogenesis, function in splicing (unwinding the U6 internal stem-loop, ISL, and annealing of U4 to U6), and recycling (Rader & Guthrie, 2002). Prp8 or Pre-mRNA-processing-splicing factor 8, one of the largest and most conserved proteins in the spliceosome, is directly implicated in splicing fidelity (Galej, Toor et al., 2018). Prp8 serves as a scaffold protein stabilizing the core of the spliceosome, which conformational rearrangement facilitates the 3D folding of the catalytically active U2/U6 RNA (Townsend, Leelaram et al., 2020).

**Figure 3.**
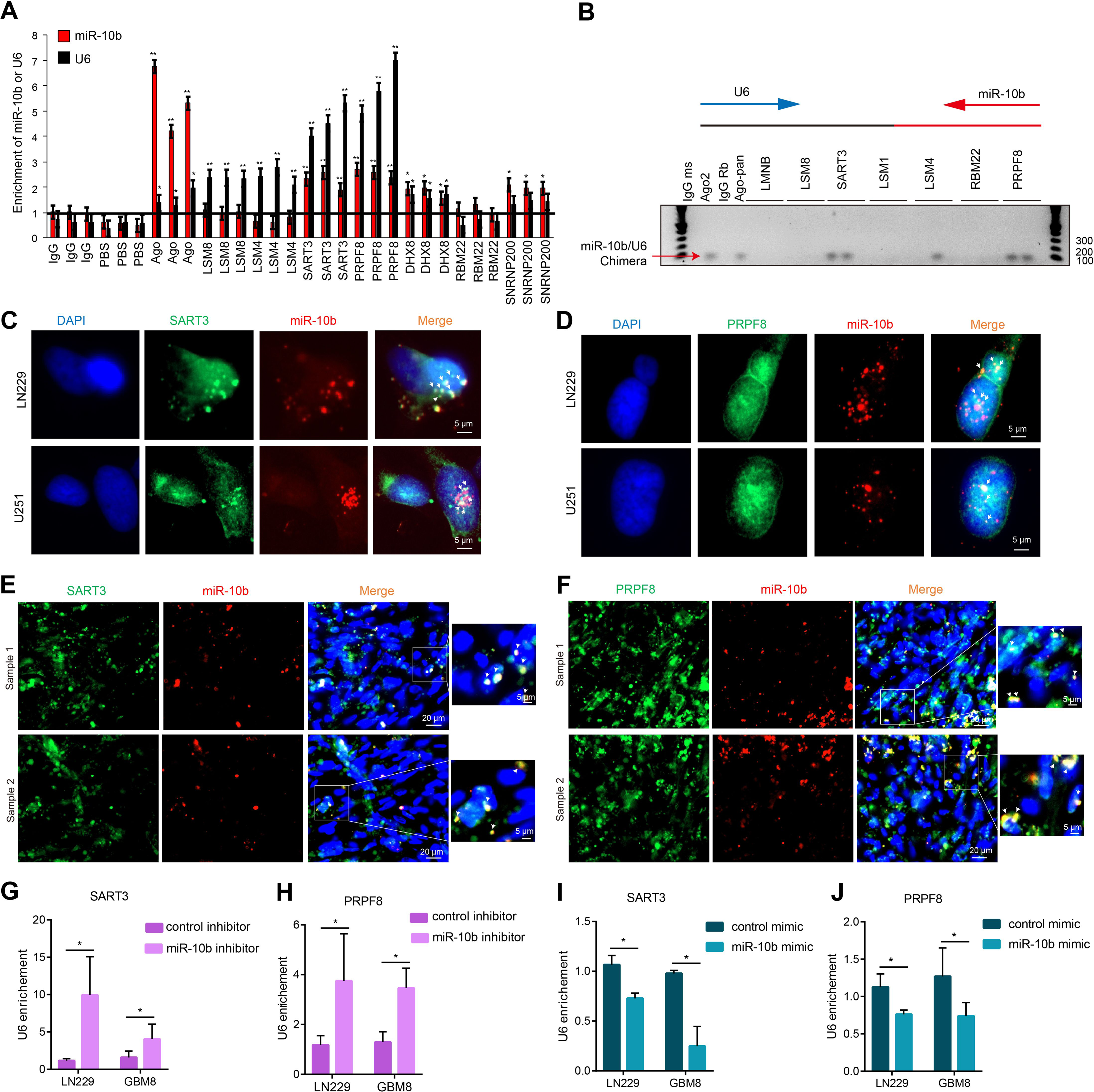
miR-10b is enriched in spliceosomal SART3 and PRPF8 RNPs. A. iCLIP of glioma cells for AGO2 and spliceosomal proteins LSM8, LSM4, SART3, PRPF8, DHX8, RBM22, and SNRNP200, followed by qRT-PCR for miR-10b and U6, demonstrates miR-10b enrichment in SART3 and PRPF8 RNPs. The results are expressed as fold-changes relative to the IgG (mean ± SD, n=3). *P* values were calculated using two-tail unpaired t-test. * *P*< 0.05; ** *P* < 0.01. B. miR-10b-U6 chimeras were detected with a pair of PCR primers, one corresponding to miR-10b and another to U6, using iCLIP assay with AGO2 and Pan-Ago antibodies, and antibodies against LMNB and splieosomal factors LSM8, SART3, LSM1, LSM4, RBM22, PRPF8 in LN229 cells. C, D. Representative images of miR-10b FISH (red) and either SART3 (C) or PRPF8 (D) immunofluorescence (green) in glioma cells, and nuclei stained with DAPI (blue) (n=3). Arrows mark the miR-10b colocalization to SART3 and PRPF8 RNPs. E, F. Representative images of the colocalization of miR-10b FISH (red) and either SART3 (E) or PRPF8 (F) immunofluorescence (green) in human GBM tumors, and nuclei stained with DAPI (blue) (n=3). Arrows mark the miR-10b colocalization to SART3 and PRPF8 RNPs. G-J. iCLIP with SART3 and PRPF8 antibodies on LN229 and GBM8 cells, transfected with either miR-10b inhibitor or mimic and the corresponding control oligonucleotides, followed by qRT-PCR detection of U6, demonstrate that U6 is displaced from the RNPs by miR-10b (mean ± SD, n=3). *P* values were calculated using two-tail unpaired t-test. * *P* < 0.05.

Using a combination of miRNA FISH and immunofluorescence (IF) techniques, we further visualized miR-10b in the nuclear SART3 speckles observed in glioma cells (Fig 3C). miR-10b also partly colocalized with PRPF8 that was distributed in the nuclei more diffusely (Fig 3D). Moreover, miR-10b colocalization with nuclear SART3 and PRPF8 RNPs was observed in patients-derived GBM tumors (Fig 3E, F). Of note, miR-10b binds to the nucleotides 79-89 within the asymmetric bulge of U6, which is the SART3 binding site, suggesting the competitive manner of miR-10b and SART3 binding to U6. To test this hypothesis, we transfected glioma cells with the miR-10b antisense oligonucleotide (ASO) inhibitor and examined the effects of miR-10b KD on the U6 association with SART3 and PRPF8 using iCLIP. U6 association with both SART3 and PRPF8 was enhanced by the miR-10b inhibitor (Fig 3G, H). Conversely, it was reduced by the synthetic miR-10b mimic (Fig 3I, J). These results indicate that miR-10b interferes with U6 incorporation into SART3 and PRPF8 snRNPs. Of note, both SART3 and PRPF8 were pulled-down by the Ago2 immunoprecipitation. Conversely, Ago2 and pan-Ago were immunoprecipitated in the SART3 and PRPF8 complexes (Fig EV3C, F). These data further support the interference of the miRNA and the splicing machineries.

### miR-10b regulates U6 snRNA levels, stability, modifications, and conformation

miRNAs are established regulators of gene expression that destabilize mRNA targets and reduce their translation. Here we investigated whether miR-10b regulates the levels of U6 snRNA. Two alternative techniques, qRT-PCR and Northern blot, demonstrated that miR-10b overexpression reduced U6 levels similarly to the effects induced by two alternative, fully complementary U6 ASOs (Fig 4A-C). Conversely, miR-10b inhibitor upregulated the U6 levels (Fig 4B, C). These effects were specific for U6 as other small RNAs were not affected by miR-10b modulators (Fig 4C and EV4A). Furthermore, the experiments with transcription inhibitor actinomycin D demonstrated that miR-10b reduced the stability of U6, with miR-10b inhibitor and mimic causing stabilization and de-stabilization of U6, respectively (Fig 4D). These effects appeared independent of the conventional RISC machinery since a luciferase reporter with the inserted full-length U6 sequence has been affected by neither miR-10b inhibitor nor its mimic (Fig EV4B).

**Figure 4.**
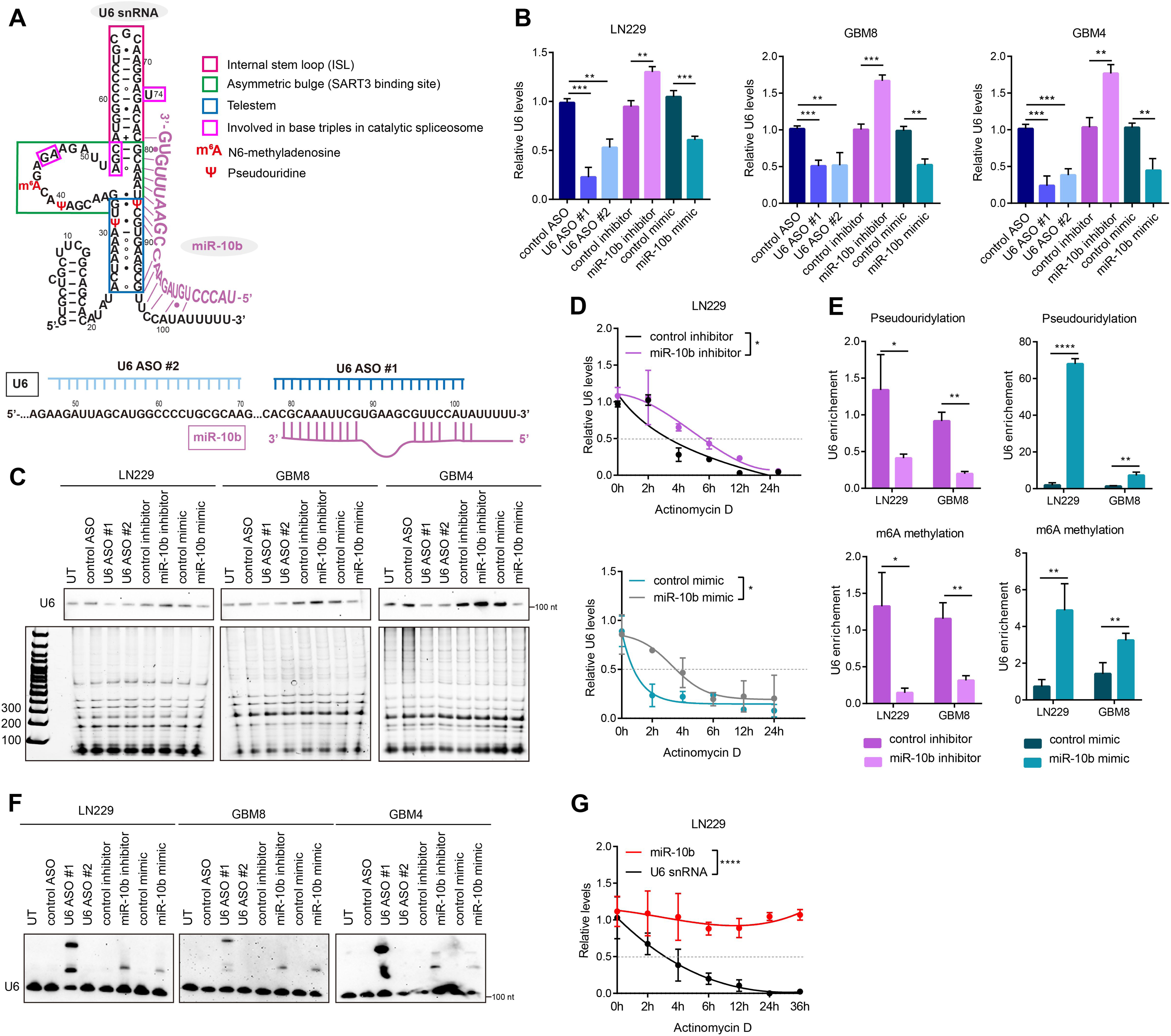
miR-10b regulates U6 snRNA levels, stability, and conformation. A. Sequence, putative secondary structure, nucleotide modifications of human U6, and miR-10b binding to U6 (top panel). The structure and modifications of human U6 have not been experimentally determined, and are shown to mimic that of yeast U6 (Didychuk et al., 2018). Positions of U6-targeting ASOs are shown in the lower panel. U6 ASO-1 binds to the 3’end of U6, similarly to miR-10b. B. qRT-PCR analysis of U6 levels in glioma cells and GSCs transfected with either U6 ASOs, miR-10b inhibitor, mimic, or corresponding control oligonucletides (mean ± SD, n=3). *P* values were calculated using two-tail unpaired t-test. ** *P* < 0.01; *** *P* < 0.001. C. Northern blot analysis of glioma cells transfected with either U6 ASOs, miR-10b inhibitor, or mimic, with a U6-specific probe, demonstrates that U6 levels are regulated by miR-10b. The lower panel demonstrates that other small RNA species, resolved by denaturing electrophoresis, are not regulated by miR-10b. D. qRT-PCR analysis of U6 levels in LN229 cells transfected with miR-10b inhibitor (top) or mimic (bottom), and treated with 5 µg/ml Actinomycin D demonstrates that miR-10b reduces U6 half-life (mean ± SD, n=3). *P* values were calculated using two-way ANOVA. * *P* < 0.05. E. iCLIP with antibodies recognizing pseudouridylation (top) and m6A methylation (bottom) on LN229 and GBM8 cells transfected with either miR-10b inhibitor or mimic, and the corresponding control oligonucleotides, followed by U6 qRT-PCR detection, demonstrate that U6 modification are regulated by miR-10b (mean ± SD, n=3). *P* values were calculated using two-tail unpaired t-test. * *P* < 0.05; ** *P* < 0.01; **** *P* < 0.0001. F. Representative native Northern blotting with U6-specific probe demonstrates that miR-10b modulation leads to the appearance of an additional U6 structural variant. G. Glioma cells treated with Actinomycin D, followed by qRT-PCR for miR-10b and U6 levels, exhibit high stability of miR-10b (mean ± SD, n=3). *P* values were calculated using two-way ANOVA. **** *P* < 0.0001.

To better understand how miR-10b regulates U6 stability, we investigated its effects on the selected U6 RNA modifications. U6 is pseudouridylated at several positions (Aoyama, Yamashita et al., 2020, Vitali, Royo et al., 2003) and miR-10b binding to the U6 nucleotides 79-88 could involve the constitutively pseudouridylated U at position 86 (U86). It may also interfere with U31 pseudouridylation of the telestem and other U6 snRNA modifications and intramolecular secondary structures (Fig 4A). Using iCLIP technique, we tested the effects of miR-10b modulation on the U6 pseudouridylation. miR-10b mimic strongly induced its levels whereas miR-10b inhibitor reduced it (Fig 4E). Similar effects on the N-6-methyladenosine modification (m6A) of U6 found at position A43 have also been observed (Fig 4E).

During the splicing cycle, U6 snRNA undergoes series of conformational changes required for accurate interactions with other snRNA and protein components of the spliceosome, recognition of mRNA splice sites and intron removal (Didychuk, Butcher et al., 2018, Nguyen, Galej et al., 2015). We hypothesized that miR-10b binding affects U6 conformation and utilized native PAGE and Northern blot hybridization to test this hypothesis. Notably, transfection of glioma cells with both miR-10b mimic or inhibitor led to the appearance of a slower-migrating U6 variant, similar to one of the variants formed in the cells transfected with U6 ASO1 that also binds to the 3’ end of the U6, as does miR-10b (Fig 4F). Altogether, these data indicate that miR-10b binding to U6 snRNA modulates U6 modifications, conformation, its binding to SART3 and PRPF8 protein partners in the snRNP assembly, and overall U6 stability.

miR-10b is much shorter than U6. Furthermore, based on glioma small RNAseq datasets, miR-10b is estimated to be ∼ 4-140-fold less abundant than U6 (Wei, Batagov et al., 2017), raising the question of how miR-10b regulates U6. In contrast to snRNAs, miRNAs are among the most stable cellular transcripts (Reichholf, Herzog et al., 2019). We, therefore, hypothesized that a single copy of miR-10b could exert its function on multiple U6 complexes. To directly compare half-lives of the endogenous miR-10b and U6 in glioma, we carried out the experiments using actinomycin D. Notably, the U6 half-life is about 3-4 hrs; however, miR-10b exhibits much higher stability, with the half-life of at least 36 hrs (Fig 4G and EV4C, D). These results suggest that a single miR-10b molecule can bind to and perturb the conformation and function of multiple U6 snRNAs during its life cycle.

### miR-10b and U6 regulate alternative splicing of CDC42

miR-10b has been previously implicated in regulating alternative splicing via targeting splicing factors such as SRSF1 and MBNL1-3 (Ramanujan, Cravens et al., 2021, Teplyuk et al., 2016). The observed direct regulation of U6, the core component of the spliceosome machinery, however, suggests that miR-10b may also have U6-mediated effects on splicing. The absolute and relative levels of U6 in the spliceosome are highly variable during development, across tissues, and cancer samples, and U6 perturbations have profound impacts on gene-specific differences in alternative splicing but not transcriptome-wide splicing failure (Dvinge, Guenthoer et al., 2019). Consistent with this, the RNA sequencing demonstrated global changes in splicing patterns upon treatments of GSCs with miR-10b inhibitor and U6 ASO (Fig 5A, Datasets EV3 and EV4). Notably, alternative splicing of numerous genes changed similarly by both miR-10b inhibitor and U6 ASO1, with similarities observed in exon skipping (SE), alternative 5′ splice sites (A5SS), alternative 3′ splice sites (A3SS), mutually exclusive exons (MXE), and retained introns (RI) (Fig 5A, Datasets EV3 and EV4).

**Figure 5.**
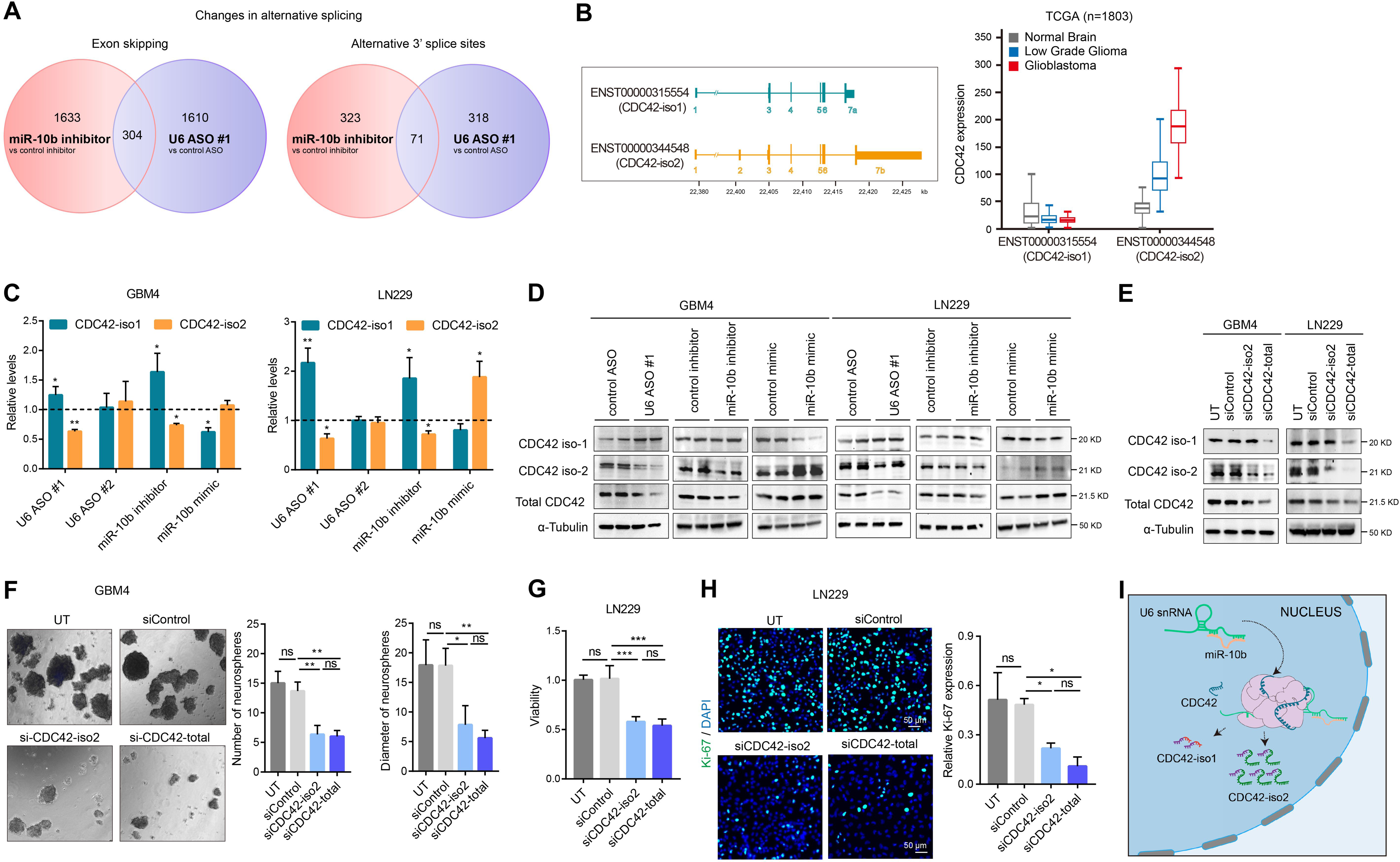
miR-10b regulates alternative splicing of CDC42, thereby controlling its total levels via U6 regulation. A. Alternative splicing events induced by miR-10b inhibitor and U6 ASO #1, as determined by RNAseq (n=3). Venn diagrams indicate the numbers of exon skipping (SE) and alternative 3’ splice sites (A3SS) modulation, in the indicated conditions. B. Schematic illustration of two major CDC42 mRNA isoforms, with alternative exons encoding alternative 5’ and 3’ UTRs (left panel). Expression analysis of CDC42 isoforms in the normal brain (n=1141), low grade glioma (LGG) (n=509) and GBM (n=153) in TCGA database demonstrates that ENST00000344548 (CDC42-iso2) is the pathologic variant associated with glioma progression, whereas ENST00000315554 (CDC42-iso1) is present at similarly low levels in the normal brain, LGG, and GBM (right panel). C. qRT-PCR analysis of CDC42-iso1 or iso2 mRNAs in glioma cells and GSCs transfected with either U6 ASOs, or miR-10b inhibitor or mimic. The results are expressed as the fold-changes relative to the corresponding control groups (mean ± SD, n=3). *P* values were calculated using two-tail unpaired t-test. * *P* < 0.05; ** *P* < 0.01. D. Western blot analysis of the indicated CDC42 forms in the cells transfected with either U6 ASO, or miR-10b inhibitor or mimic. E. Western blot analysis of the cells transfected with either selective siRNA targeting iso2 (siCDC42-iso2) or both CDC42 isoforms (siCDC42-total) demonstrates that KD of iso2 is sufficient for reducing total CDC42 levels. F. Analysis of GSC spheroids transfected with either siCDC42-iso2 or siCDC42-total demonstrates that CDC42 KD reduces GSC growth. The number and size of GSC spheres have been monitored using Image J at day 7 after transfection (mean ± SD, n=3, 8 images per cultured analyzed). *P* values were calculated using two-tail unpaired t-test. * *P* < 0.05; ** *P* < 0.01; ns, no significance. G. Analysis of glioma LN229 cells transfected with either siCDC42-iso2 or siCDC42-total demonstrates that CDC42 KD reduces glioma growth. Cell viability was monitored 72 hours after transfection by WST-1 assay (mean ± SD, n=3). *P* values were calculated using two-tail unpaired t-test. *** *P* < 0.001; ns, no significance. H. Representative images of LN229 cells immuno-stained for Ki-67 (green) and DAPI staining (blue) and quantification of the Ki67^+^ cells (mean ± SD, n=3). *P* values were calculated using two-tail unpaired t-test. * *P* < 0.05; ns, no significance. I. Schematic illustration of alternative splicing of CDC42, regulated by miR-10b via its binding to U6 snRNA.

To compare the effects of miR-10b and U6 perturbations, we utilized CDC42, one of the critical genes implicated in glioma biology, whose splicing is modulated by miR-10b (Teplyuk et al., 2016). There are two major CDC42 mRNA variants produced in GBM: the isoform ENST00000315554 (iso-1) and the predominant variant ENST00000344548 (iso-2) with the additional exon 2 and alternative last exon utilized (Fig 5B). Although the ORFs of the two isoforms are very similar and translate to the proteins with only slightly altered C-terminus, their 3’ and 5’ UTRs are highly distinct. Iso-2 mRNA is significantly more stable than iso-1 (Fig EV5A), consistent with its prevalence in glioma. Modulation of miR-10b by its inhibitor and mimic had reciprocal effects on the levels of two isoforms, with iso-1 being downregulated and iso-2 upregulated by miR-10b and resulting in the altered total levels of CDC42 protein (Fig 5C, D and EV5B, C). Notably, the ASO complementary to 3’ end of U6 snRNA (ASO1) and highly similar to miR-10b, but not another ASO (ASO2), modulated CDC42 splicing and overall expression, similarly to the miR-10b inhibitor (Fig 5C, D and EV5B, C).

miR-10b inhibition reduces glioma growth and leads to glioma cell death *in vitro* and in animal models *in vivo* (Gabriely et al., 2011b, Lin, Teo et al., 2012, Teplyuk et al., 2016). CDC42 small Rho GTPase has also been implicated in cancer cell proliferation and viability (del Mar Maldonado & Dharmawardhane, 2018). We, therefore, questioned whether CDC42 could partly mediate miR-10b effects on glioma growth. Downregulation of CDC42 by siRNAs, either cognate to the iso-2 or the common region of the iso-1 and iso-2 transcripts, to the levels mimicking the effect of the miR-10b inhibitor (Fig 5E and EV5D, E), significantly reduced glioma proliferation and growth, as indicated by Ki67 staining, number and size of GSC spheroids, and overall cell viability (Fig 5F-H and EV5F). We, therefore, concluded that miR-10b exerts its glioma-promoting activity, in part, by binding U6 snRNA and thereby affecting CDC42 splicing (Fig 5I). This, in turn, leads to the expression of the more stable CDC42 mRNA isoform and increased protein levels of this important growth-promoting factor.

## DISCUSSION

MiR-10b has been established as an onco-miR associated with various malignancies (Lund, 2010, Tehler et al., 2011). It is most strongly upregulated in malignant gliomas and required for glioma and GSC viability (Gabriely et al., 2011b, Teplyuk et al., 2016). Nevertheless, there is still little understanding of the striking glioma “addiction” to miR-10b and its signaling in cancer more generally. Even for a well-defined malignancy such as the GBM, the attempts to study molecular mechanisms underlying miR-10b function are hampered by the (a) inter- and intratumoral heterogeneity of the tumors and multiple mechanisms of miR-10b activity in genetically diverse glioma cells, and (b) unconventional targeting properties of this miRNA (Teplyuk et al., 2016). Previous studies suggested that miR-10b commonly binds and regulates target mRNAs by a non-canonical, non-seed-mediated mechanism, and control pre-mRNA alternative splicing (Teplyuk et al., 2016). In this study we employed a recently developed CLEAR-CLIP technology for direct transcriptome-wide capture of miR-10b interactions and identification of its targets in glioma. Since the technique is based on the cross-linking and ligation of the spatially proximal transcripts, it may reveal both direct and more complex indirect interactions between the transcripts, including false-positives (Moore et al., 2015). Initially proposed quality metrics for CLEAR-CLIP relied on the bioinformatical analysis demonstrating that miRNA-mRNA chimeras are enriched in a miRNA seed. However, such approach is unconstructive for miRNAs that operate via non-canonical binding. Therefore, although our analysis identified more than a hundred of putative targets with miR-10b binding sites (mostly seedless, see below), additional experimental work is required for their functional validation. Here we used several complementary approaches to validate an unanticipated top miR-10b target-a short regulatory snRNA U6, rather than a conventional mRNA. Despite the significant basepairing and abundant binding to miR-10b, U6 was not predicted by current algorithms developed with the focus on mRNA and further updated to include long non-protein-coding RNA species. miR-10b-U6 interactions have been confirmed biochemically, with chimeras observed in AGO2, pan-AGO, as well as snRNP (SART3, PRPF8) immunoprecipitates. Enrichment of miR-10b in high-density nuclear fractions resolved in glycerol gradient, containing pre-catalytic spliceosome, further supports these results. Furthermore, small RNA FISH that we developed for miR-10b detection revealed its co-localization with U6 snRNA and U6 snRNPs in glioma cells and patient-derived tumors. Finally, the chimeric reads produced by miR-10b and U2 snRNA, another spliceosomal RNA and U6 -binding partner, further substantiated miR-10b interaction with the spliceosomal machinery.

U snRNAs are essential components of the spliceosome, required for the recognition of substrate pre-mRNAs and serving as ribozyme catalysts of two consecutive transesterification reactions to ligate two exons, concurrent with an intron removal (Wahl, Will et al., 2009). Cryo-EM structures demonstrated that U6 snRNA forms the catalytic core of the spliceosome and interacts with three other snRNAs (U2, U4, and U5), >25 protein partners, and the pre-mRNA substrate throughout the splicing cycle, which is accompanied by its large conformational changes (Didychuk et al., 2018, Wahl et al., 2009). MiR-10b binds to a highly conserved and functionally important part of the U6 at its 3’ end and is anticipated to interfere with the formation of major U6 structures, including internal stem loop (ISL), telestem, and asymmetric bulge (a SART3 binding site) (Fig 4A). These secondary structures undergo dynamic transitions into other conformations during the spliceosome assembly and activation. For example, incorporation of U6 into the precatalytic spliceosomal B complex (one of the key steps of the spliceosome assembly) requires unwinding of the ISL and subsequent annealing to U4 snRNA (Didychuk et al., 2018). The protein SART3 is essential for this reaction (Didychuk et al., 2018, Rader & Guthrie, 2002). As we demonstrate, it can be displaced from U6 by miR-10b. miR-10b also interacts with PRPF8, a scaffolding factor that mediates the orderly assembly of proteins and snRNAs in the pre-catalytic U4/U6-U5 snRNP complex. The U6 telestem, another miR-10b-interacting structure, is mutually exclusive with the formation of U2/U6 helix II required for the spliceosome activation (Ast, Pavelitz et al., 2001, Sashital, Cornilescu et al., 2004). Therefore, since U6 secondary and tertiary structures formed in this region are critically important for the spliceosome function and miR-10b modulates these structures, it has a major impact on this fundamental cellular machinery. miR-10b binding may also interfere with essential nucleotides in the U6, such as the highly conserved catalytic AGC triad that is critical for splicing catalysis, and the adjacent ACAGAGA box that interacts with the 5′ splice site of pre-mRNA and helps organize the spliceosome active site through the formation of base triple interactions (Fig 4A) (Fica, Mefford et al., 2014).

MiR-10b also promotes U6 post-transcriptional modifications, including pseudouridylation and m^6^A modification. m^6^A, the most prevalent internal modification associated with eukaryotic RNAs, influences many steps of mRNA metabolism and may be important for regulatory functions of ncRNAs as well. In U6 snRNA, a single m^6^A at position A43 (U6–A43) is modified by a recently identified methyltransferase, METTL16 (Pendleton, Chen et al., 2017, Warda, Kretschmer et al., 2017). METTL16 activity on U6–A43 depends on the formation of the adjacent ISL secondary structure within U6 (Shimba, Bokar et al., 1995). While m^6^A-containing mRNAs undergo distinct pathways of rapid degradation (Lee, Choe et al., 2020), whether and how m^6^A regulates U6 function and stability in human cells is unknown. Interestingly, the recent yeast-based study demonstrated global splicing alterations in a strain lacking U6–A43 modification and suggested a function for this modification in 5’splice site recognition and exon diversification (Ishigami, Ohira et al., 2021). This is consistent with the A43 positioned in the center of the ACAGAGA motif essential for U6 base pairing to the intron adjacent to the 5′ splice site of a pre-mRNA (Fica et al., 2014).

Human U6 is also constitutively pseudouridylated at several positions (Massenet & Branlant, 1999), including U86 that could directly participate in the base-pairing with miR-10b, and U31 and U40, positioned in the telestem and SART3-binding region; they could also be affected by miR-10b and miR-10b-modulated SART3 binding. The effects of these modifications on U6 structure, stability, and functions are largely unknown, although studies of *Saccharomyces cerevisiae* suggest their functionality (Basak & Query, 2014). Pseudouridylation is generally thought to be structurally stabilizing because of the high base stacking (Borchardt, Martinez et al., 2020); however, this notion stems from mRNA studies and may be irrelevant for the highly structured small RNA such as U6. Our study demonstrates that miR-10b increases the levels of U6 pseudouridylation and m^6^A modification, but downregulates the overall U6 stability, suggesting the negative effects of the modifications on U6 stability. This could be due to the highly structured character of U6 and binding to multiple RBPs, the characteristics affected by the modifications.

Altogether, our data suggest the mechanistic relationships between miR-10b binding to U6 and its effects on U6 modifications, conformation, and destabilization. However, the molecular mechanisms and the sequence of events remain to be further investigated. We propose that miR-10b binding leads to a U6 structural-conformational reorganization that, in turn, alters protein binding of some snRNP proteins (e.g., SART3, PRPF8) and enzymes catalyzing U6 modification, including writers and erasers (e.g., methyltransferase METTL16 and pseudouridine synthase Pus1). This, in turn, regulates U6 modifications, spliceosome assembly, and splicing reactions. In such a scenario, the lowered levels of U6 could be a consequence of the reduced U6 incorporation in the snRNPs. Alternatively, miR-10b binding may first facilitate the recruitment of U6 modifiers leading to the increased levels of modifications and thereby affecting the U6 structure, steady-state, and splicing cycle. miR-10b binding at the 3’end of U6 can also directly interfere with U6 trimming and turnover. Regardless of the order of events, miR-10b-mediated effects on U6 secondary structure, which is tightly controlled during the splicing cycle, must play a critical role in the downstream effects on mRNA splicing. The data indicating that miR-10b inhibitor and U6 ASO1 that cause similar conformation changes in U6, but reciprocal effects on U6 levels, have nearly identical effects on the CDC42 splicing (and splicing patterns of numerous other genes), further support the importance of miR-10b-mediated U6 reorganization, rather than absolute U6 levels, in splicing catalysis. In part, the structural alterations may result from miR-10b displacement of major U6 binding proteins, such as SART3. Intriguingly, in addition to displacing SART3 from U6, miR-10b also directly reduces SART3 expression by targeting its mRNA (Teplyuk et al., 2016). Such a two-fold effect on SART3 may provide a mechanism of enhanced control over U6 structure and function.

One of the important questions regarding miR-10b regulation of U6 snRNA relates to the stoichiometry of the two molecules. U6 is an abundant transcript, which level reaches 100,000 copies/ cell in HeLa cells. Many miRNAs also accumulate to very high steady-state levels, with some at least as plentiful as the U6 snRNA (Lim, Lau et al., 2003). Our analysis of glioma datasets (Wei et al., 2017) suggested that in diverse GSC subtypes the ratio of miR-10b/ U6 ranges from 1:140 to 1:4. How can miR-10b control the structure and functions of U6 that is present at much higher levels? The experiments with Actinomycin D indicate that miR-10b is about 10-times more stable than U6. Furthermore, since it is by far more potent inhibitor of RNA Pol II that transcribes miR-10b (active at 0.5 µg/ml) than of Pol III transcribing U6 (5 µg/ml) (Bensaude, 2011), our experiments likely underestimate the difference in the stability of the two molecules. The highly stable miR-10b, therefore, may bind to and regulate U6 via a catalytic mechanism, with a single miR-10b copy capable of interfering with multiple U6 RNPs.

Furthermore, as miRNA binding to its targets is one of the factors defining the miRNA half-life (Ghini, Rubolino et al., 2018), ample binding to U6 and U6 RNPs may contribute to the miR-10b steady-state levels.

Our work contributes to the growing body of evidence supporting miRNA nuclear localization and non-canonical functions and expands the mechanisms and repertoire of miRNA targets. In contrast to a canonical pre-miRNA undergoing nuclear export, cleavage by the RNase III Dicer, and loading of a mature miRNA to the RNA-induced silencing complex (RISC) in the cytosol, where it targets mRNAs, some mature miRNAs are found in the nucleus. Among the first reported nuclear miRNAs, for example, is miR-21-another important onco-miR implicated in the carcinogenesis (Meister, Landthaler et al., 2004). We demonstrate that about 50% of miR-10b in glioma cells is nuclear, but the subcellular trafficking and selectivity of such localization remain to be further investigated. miR-10b could be either produced by an atypical, entirely nuclear biogenetic pathway or undergo the cytosolic maturation followed by the reverse cytosol-to-nucleus shuttling. Nuclear localization of miRNAs can be mediated by different sequence motifs (e.g., AGUGUU or UUGCAUAGU), in a mechanism controlled by specific RBPs. However, miR-10b does not contain such nuclear localization motifs. At least two pieces of evidence provide support for the nuclear miR-10b maturation scenario: first, the finding of Ago2 and, more recently, of Dicer in the nucleus of specific cells, including glioma (Bronisz et al., 2020, Burger, Schlackow et al., 2017, Castanotto, Zhang et al., 2018, Chu, Yokota et al., 2021, Gagnon, Li et al., 2014, Liu et al., 2018), and second, the co-IP of the spliceosomal SART3 and PRPF8 complexes with Ago2, the core endonucleolytic components of RISC that efficiently pools-down the entire miR-10b population from glioma cells. There is also emerging evidence that RISC components continuously shuttle between the nucleus and cytoplasm (Liu et al., 2018). Although prior data indicate that miRNA-Ago complexes target nuclear transcripts, including mRNAs and pre-mRNAs (Bottini, Hamouda-Tekaya et al., 2017, Chu et al., 2021, Sarshad, Juan et al., 2018), here we characterize the first miRNA target that is a small regulatory RNA essential for eukaryotic cells. This study, therefore, expands the miRNA-regulated networks beyond mRNA and lncRNA targets and suggests exciting new avenues for miRNA research.

In addition to discovering snRNA as a new class of miRNA targets, our work indicates that an established oncomiR-10b targets multiple transcripts in a noncanonical way, in most cases based on the non-seed and non-3’UTR binding. Although the seed sequences confer the strongest mRNA degradation, functional seedless sites have been discovered in multiple biologically important mRNAs (Bjerke & Yi, 2020, Chipman & Pasquinelli, 2019, Flamand, Gan et al., 2017, Moore et al., 2015, Yang, Bofill-De Ros et al., 2019, Zhang et al., 2015). For example, a new type of miRNA-recognition elements found exclusively in CDSs depend on miRNA 3’-end interactions, but not seed pairing. This type of binding blocks translation elongation in an Argonaute-dependent but GW182-independent manner, resulting in reduced protein but not mRNA levels of the targets (Zhang, Zhang et al., 2018). This finding is consistent with our results demonstrating that the majority of seedless miR-10b interactions are found in the CDSs. Of note, some developmentally essential miRNAs, such as miR-9, appear to favor seedless interactions (Moore et al., 2015). Furthermore, we demonstrate that seedless interactions can interfere with RBP binding and modulate target’s structure, thereby affecting its properties, and regardless of the effects on the target stability. Such miRNA control could be especially important for altering structural characteristics and activity of regulatory transcripts, including small and long ncRNAs. Investigation of the normal physiological and pathology-associated networks formed by interacting miRNAs and other regulatory RNA species represents an exciting new avenue for fundamental and translational research.

## MATERIALS AND METHODS

### Cell culture

LN229, U251 (human glioblastoma), MDA-MB-231 (human breast cancer) and HEK-293 (human embryonic kidney) and Hela cell lines cultured in Dulbecco’s Modified Eagle Media (DMEM) supplemented with 10% Fetal Bovine Serum (FBS) (WISENT), 1% penicillin and streptomycin (Gibco) and passaged by trypsinization (Corning^®^). SH-Sy-5y (human neuroblastoma) cell line was purchased from ATCC and passaged in DMEM-F12 supplemented with 10% FBS, 1% MEM Non-Essential Amino Acids Solution (NEAA) (Gibco), 1% penicillin and streptomycin (Gibco). Low-passage GSC (GBM4 and GBM8) cells were provided by Dr. Hiroaki Wakimoto (MGH) and cultured as neurospheres in Neurobasal medium (Gibco) supplemented with 0.5% N-2 (Gibco), 2% B-27 (Gibco), 1.5% glutamine (Gibco), 0.02 ng/ml epidermal growth factor (EGF) (Gibco), and 20 ng/ml fibroblast growth factor (FGF) (PeproTech) in ultralow attachment flasks or plates. GSC neurospheres were passaged using the NeuroCult chemical dissociation kit (Stemcell Technologies). All cells were maintained at 37 °C and 5% CO_2_.

### Oligonucleotides, siRNAs, and cell transfection

A chemically modify 2’-O-methoxyethyl ASO inhibitor of miR-10b (5’-CACAAATTCGGTTCTACAGGGTA-3’, miR-10b ASO) and control oligonucleotide of the same chemistry (5′ -ACATACTCCTTTCTCAGAGTCCA-3′) were designed as previously reported (Teplyuk et al., 2016). The following U6 ASOs and corresponding control have been designed based on the (Dvinge et al., 2019, Liang, Vickers et al., 2011): U6 ASO#1 mUmGmGmAmACGCTTCACGAATmUmUmGmCmG, U6 ASO#2 mUmGmAmUmAAGAACAGATACTACAmCmUmUmGmA; control mCmCmAmUmGGCTAACCACAmUmUmCmUmU (m=2’-O-methyl, with phosphorothioate backbone). MiR-10b mimic (cat no: C-300550-07-0005) and matching control (cat no: CN-001000-01-05), CDC42-isoform2-specific siRNA (AGCCUCCAGAACCGAAGAAGA) and control siRNA (UGGUUUACAUGUCGACUAA), as well as all oligonucleotides listed above were obtained from Dharmacon. CDC42-total siRNA was a pool of 4 siRNAs targeting both isoforms (CUCCUGAUAUCCUACACAA; CCUAGAUCUAGUUUAGAAA; GAGUAGAUCCAGUAUUUGA; CCCAAACCUAAUUCUUGUA), purchased from Santa Cruz Biotechnology (sc-29256). For transfections, adherent LN229 and U251 cells were plated in 24 well plates (Corning ^®^) at 2× 10^4^/ well. On the next day, the cells were transfected with either miR-10b inhibitors or mimics (final concentration of 50 nM), U6 ASOs (final concentration of 1 nM), CDC42 siRNAs (final concentration of 100 nM), or corresponding control oligonucleotides using lipofectamine 2000 (ThermoFisher Scientific). For GSC transfection, neurospheres were dissociated to single cell suspension (Stemcell Technologies) and transfected with the oligonucleotides using Neuromag 500 (OZ Biosciences).

### RNA isolation and Real-time quantitative PCR (qRT-PCR)

RNA was extracted using the Total RNA Purification Kit (Norgen Biotek, Catalogue # 17250) and quality of the samples examined using Nanodrop. For microRNA analysis, 200 ng of the total RNA was reverse transcribed using miRCURY LNA RT kit (QIAGEN, 339340). The cDNA libraries were then diluted 80 times and used for the analysis of gene expression by qPCR using miRCURY LNA SYBR Green PCR kit (QIAGEN, 339347) with miRCURY LNA™ primers (YP00203907 for U6; YP00205637 for miR-10b, YP00204339 for miR-125a) (QIAGEN). For mRNA analysis, 200 ng of the total RNA was reverse transcribed to cDNA (BIO-RAD, #1708841). The qPCR was run on Viia 7 RT-PCR system (Applied Biosystems). The fold-changes in gene expression were calculated by ΔΔCt method. The expression of miRNA and mRNA was normalized by miR-125a and GAPDH, respectively. The following primers have been used: CDC42 isoform1: F 5’-GCCTATCACTCCAGAGACTGC-3’, R 5’-GCTCGAGGGCAGCTAGGATA-3’; CDC42-isoform2: F 5’-GCCTATCACTCCAGAGACTGC-3’, R 5’-GCTCCAGGGCAGCCAATATT-3’; GAPDH: F 5’-ATGTTCGTCATGGGTGTGAA-3’, R 5’-TGTGGTCATGAGTCCTTCCA-3’.

### Denaturing and Native Northern blot

RNA samples were analyzed by denaturing Northern blotting, following the manufacturer’s instructions (Signosis, High Sensitive MiRNA Northern Blot Assay Kit, NB1001). Briefly, total RNA (300 ng) was denatured at 70°C for 5 min and resolved on 4-20% Tris-borate-EDTA (TBE) Urea gel (ThermoFisher scientific) in cold 0.5 × TBE buffer (Invitrogen) at 60V for 5-6 hrs. The RNA was then transferred to a nylon membrane at 60V in 0.5× TBE with 20% methanol for 1 hr at 4 °C. The RNA was crosslinked to the membrane by UV irradiation (120mJ/cm²). The membranes were incubated with biotin-pre-labeled probes overnight at 42°C and rinsed with hybridization washing buffer for 30min at 42°C. After blocking, the membranes were incubated with Streptavidin-HRP conjugate and signals developed using a chemiluminescence imaging system. The oligonucleotide probes for Northern blots were as following: U2 (ACACTTGATCTTAGCCAAAAGGCCGAGAAGCGAT); U4 (GCCAATGCCGACTATATTGC); U6 (AGTATATGTGCTGCCGAAGCGAGCACT). For the analysis of native RNA conformation, cells were washed and resuspended in a mild lysis buffer (1% lgepal, 0.5% sodium deoxycholate, 0.1% SDS), and deproteinized in a buffer containing 1 mg/ml proteinase K, 50 mM NaCl, 10 mM EDTA, and 0.5% SDS for 30 min at 37°C. Total RNA (5 μg) was separated in 4-20% TBE gel (ThermoFisher scientific), described above.

### RNA stability and half-life

For monitoring half-life of miR-10b and U6 snRNA, glioma LN229 and U251 cells were cultured in 6-well plates. The cells were treated with Actinomycin D (final concentration 5 µg/ml, in 0.05% DMSO) (Sigma) for 2-36 hrs, followed by the RNA isolation and analysis. To examine the effects of miR-10b on U6 stability, cells were transfected with either miR-10b inhibitor or mimic and matching control oligonucleotides and culture media was replaced with the media containing 5 μg/ml of Actinomycin D at 24 hours after transfections. RNA was isolated at different time points and qRT-PCR reactions were performed as described above.

### Western blot

Cells were harvested using RIPA buffer (Boston BioProducts) and proteins were quantified using Micro BCA^TM^ Protein assay kit (ThermoFisher Scientific). The cell lysates containing equal amounts of total proteins (30 μg) were separated by SDS PAGE in 4-12% Bolt Bis-Tris Plus gels (ThermoFischer Scientific) and transferred to 0.2 µm Immun-Blot^®^ PVDF membrane (BIO-RAD). After blocking with 5% (w/v) fat free milk in phosphate buffer saline (PBS) with 0.1% Tween-20 (PBS-T), membranes were incubated with primary antibodies (anti-HSP90 (Cell Signaling Technology, #4874S, 1:3000 dilution); anti-Lamin B (ABclonal, A16685, 1:1000 dilution); anti-CDC42-iso1 (Millipore, ABN1646, 1:1000 dilution); anti-CDC42-iso2 (07-1466, Millipore, 07-1466, 1:1000 dilution); anti-CDC42-total (PA1-092, ThermoFischer Scientific, PA1-092, 1:1000 dilution); α-Tubulin (Sigma, T9026, 1:500 dilution) at 4 °C overnight. The membranes were further washed with PBS-T buffer and incubated with secondary antibodies (Cell Signaling Technology, #14708 or #14709, 1:10,000 dilution) for 2 hrs at room temperature. The blots were developed using ECL reagent Supersignal West Pico chemiluminescent Substrate (ThermoFisher Scientific).

### Cell fractionation and isolation of nuclear fractions

For subcellular fraction, 5 × 10^6^ cells were collected and washed twice with PBS and transferred to a microcentrifuge tube. The pellets were re-suspended in 1x hypotonic buffer (20 mM Tris-HCl (pH= 7.4), 10 mM NaCl, 3 mM MaCl_2_) and incubated on ice for 5 min. 1% of a Non-Ionic Detergent (NP40) was added and the lysates gently pipetted and incubated for 5 min, followed by centrifugation of the homogenates for 10 min at 3,000 rpm at 4 °C. The supernatants containing cytosolic fractions and the pellets containing nuclear fractions have been collected. The pellets were resuspended in the cell extraction buffer (10 mM Tris (pH= 7.4), 2 mM Na_3_VO_4_, 100 mM NaCl, 1% Triton X-100, 1 mM EDTA, 10% glycerol, 1 mM EGTA, 0.1% SDS, 1 mM NaF, 0.5% deoxycholate, 20 mM Na_4_P_2_O_7_) supplemented with 1 mM PMSF and Protease Inhibitor Cocktail (Sigma), vortexed, and centrifuged again. The resulting supernatants contained soluble nuclear fractions and the pellets contained nuclear insoluble fractions. The isolated nuclear fractions were subjected to ultra-centrifugation on the linear glycerol gradient (from 5% to 60%) for 2 hrs at 4°C, 26,000 rpm using SW55 rotor (Beckman Coulter L-90K optima), and aliquoted to 200 µl fractions (Hinterberger, Pettersson et al., 1983). These fractions have been used for RNA and protein isolation and analysis.

### Immunofluorescence

Cells were washed with PBS and fixed with 4% paraformaldehyde (PFA), permeabilized with 0.5% Triton X-100 in PBS, and blocked in 5% BSA for 1 hr. The cells were then incubated with primary antibodies (rabbit anti-KI-67, Abcam, ab16667, 1:250 dilution) overnight at 4 °C and further incubated with secondary antibodies (goat anti-rabbit Alexa fluoro 488, Invitrogen, 1:200 dilution). The nuclei were stained with DAPI (1:1000). The staining was visualized and imaged using a scanning confocal microscopy.

### Fluorescence In-situ hybridization (FISH) and Immunofluorescence

RNA FISH was performed using a ViewRNA cell Plus Assay Kit (Invitrogen, #88-1900-99) following the manufacturer instructions. Briefly, 1×10^4^ cells per well were cultured in a chamber slide (8 Chamber Polystyrene Vessel Tissue Culture Treated Glass Slide, FALCON, #354118) and fixed with the fixation solution for 1 hr at room temperature. For the tumor analysis, resected GBM specimens were obtained from the Brigham and Women’s Hospital, per a protocol approved by the Institutional Review Board. Frozen GBM tissues were fixed with 4% PFA overnight at 4 °C and blocked in 30% sucrose for 4-6 hrs. After transferring to the optimal cutting temperature compound (OCT), GBM tissues were cut into 15 µm-thick sections and placed into acetone for 5 min at -20 °C. After three washes with RNase inhibitor-containing PBS, the cells/ tissue slides were permeabilized with 0.3% Triton-X 100 in PBS for 5 min on ice. The cells/tissue slides were hybridized with probes specific for mature miR-10b, miR-21, or U6 RNA (Invitrogen, 1:25 dilution) for 2 hrs at 40 °C. To control for background staining, the probes were omitted in parallel experiments. The cells were further washed with wash buffer; the tissue slides were washed with 2X SSC for 5 times at 40 °C. The cells/slides were hybridized sequentially with PreAmplifer, Amplifer and Label probe solution for 1 hr at 40 °C, stained with 1 × DAPI (4’,6-diamidino-2-phenylindole), mounted in a mounting media (ThermoFisher Scientific), and observed under a confocal microscope (CZ Scan). For the FISH combined with Immunofluorescence, cells were fixed in 4% PFA, treated with 0.3% Triton-X 100 solution, and blocked with 5% BSA. Tissues slides were placed into acetone and then treated with antigen repair solution (20 mg/ml Proteinase K, 1 M Tris-HCl (pH=8.0), 0.5 M EDTA (pH=8.0)) for 5 min at 37 °C. The step of overnight incubation at 4 °C with primary antibody (anti-PRPF8, ABclonal, A6053, 1:200 dilution; or anti-SART3, GeneTex, GTX107684, 1:200 dilution) was included before hybridization with FISH probes and the subsequent signal amplification steps. The probes were then washed and the cells/slides incubated with secondary antibodies (goat anti-rabbit Alexa fluoro 488, Invitrogen, 1:200 dilution) for 1hr at room temperature. They were further washed with PBS three times, stained with 1 × DAPI, and images taken using confocal microscopy. Quantification of the colocalization was determined by Image J pro 6.0.

### CLEAR-CLIP: Covalent Ligation of Endogenous Argonaute bound RNA -Crosslinking and immunoprecipitation

CLEAR-CLIP was performed according to the published protocol with minor modifications. Briefly, LN229 cells were irradiated with UV-C light (2 times by 2400 J/m^2^) to covalently cross-link proteins to nucleic acids. The cells were lysed, treated with DNase (Promega), followed by the partial RNA fragmentation using low concentrations of the RNase I (0.002 U/ml, 5 min), and treatment with the RNase inhibitor (RNAsin Plus at 0.5 U/μl) to quench RNase activity. The miRNA–RNA complexes were immunopurified using the anti-Ago-2 antibody (Millipore, #03-110) immobilized on immunoglobulin G-coated magnetic beads (Life Technologies). The beads were washed, treated with T4 Polynucleotide Kinase (PNK, with no 3’-phosphatase activity) (NEB, M0236S), and miRNA-RNA chimeras were produced with T4 RNA Ligase I (NEB, M0204S). The beads were further washed and subjected to alkaline phosphatase treatment (Roche, #10713023001) to remove 3’ phosphate. The 3’ linker (5′-rAppGTGTCAGTCACTTCCAGCGG-3’, Dharmacon) was added using T4 RNA Ligase 2 (NEB, M0242S), followed by the PNK (NEB, M0201S) treatment. The samples were resolved on 8% acrylamide gel (ThermoFisher Scientific), transferred to nitrocellulose membrane, followed by the purification of RNA-Ago complex, RNA isolation, 5’ linker ligation (5’-OH AGGGAGGACGAUGCGG3’-OH) (Dharmacon), and library preparation. The libraries have been prepared from the low amount of RNA input using Superscript Reserve transcriptase (ThermoFisher) with first PCR amplification performed with DP3 (5’-CCGCTGGAAGTGACTGACAC-3’) and DP5 (5’-AGGGAGGACGATGCGG-3’) primers, and additional amplification with DSFP3 (5’-CAAGCAGAAGACGGCATACGACCGCTGGAAGTGACTGACAC-3’, DSFP5 (5’-AATGATACGGCGACCACCGACTATGGATACTTAGTCAGGGAGGACGATGCGG-3’) at maximum of 16 PCR cycles. The miR-10b chimeras have been pre-amplified with a miR-10b primer TACCCTGTAGAACCGAATTTGTG and either DP3 or DP5 primers. The products were separated from primers and primer dimers using electrophoresis on a 1% agarose gel, recovered using a QIAquick gel extraction kit (Qiagen), and sequencing using llumina 150 - nucleotide paired-end multiplex sequencing at the MGH sequencing facility. For validation of miR-10b/ U6 chimeras, cDNA was amplified with a pair of PCR primers, one corresponding to miR-10b (either TACCCTGTAGAACCGAATTTGTG or CACAAATTCGGTTCTACAGGGTA) and another to U6 (GTGCTCGCTTCGGCAGCACATATAC or GTCATCCTTGCGCAGGGGCCATGC).

For identification of miRNA-RNA chimeras, 3’ and 5’ adapter sequences have been removed from reads using Trimmomatic software version 0.4.0 (Bolger, Lohse et al., 2014).The trimmed sequences were collapsed with exact duplicates removal, and mapped to the hg19 reference transcriptome with BWA-MEM aligner BWA-0.7.17 (Li, 2013). The following mapping parameters were used: seed length for 1st mapping iteration (MI) - 12; seed length for 2nd MI - 16; minimal alignment score for the 1st MI - 18; minimal alignment score for the 2nd MI - 16; matching score (MS) for 1st MI is 1; mismatch penalty (MP) for 1st MI is 4; MS for 2nd MI is 1; MP for 2nd MI is 6; gap opening penalty (GOP) for 1st MI is 6; gap extension penalty (GEP) for 1st MI is 1; GOP for 2nd MI is 100; GEP for 2nd MI is 100; maximum number of allowed multi-hits per read in 1st MI is 50; maximum number of allowed multi-hits per read in 2nd MI is 100. Chimeric reads were extracted, and miRNA/ RNA segments were identified with ChiRA workflow version 1.3.3 (Videm, Kumar et al., 2021).

### Crosslinking and immunoprecipitation (iCLIP)

To covalently crosslink the proteins to nucleic acids, glioma cells (2 × 10^7^) were subjected to UV irradiation (200 mJ/cm²). The cells were then lysed with RIPA lysis buffer containing Protease Inhibition Cocktail and RNase Inhibitor followed by DNase treatment (Promega). The immunoglobin-coated protein magnetic beads (Thermofisher scientific) were incubated with antibodies (anti-pseudouridine, MBL, D347-3; anti-N6-methyladenosine (m6A), Millipore, MABE1006; anti-PRPF8, ABclonal, A6053; anti-SART3, GeneTex, GTX107684; anti-Ago-pan, Millipore, MABE56; anti-Ago2, Millipore, 03-110; anti-LSM8, Invitrogen, PA5-69098; anti-LSM4, ABclonal, A5891; anti-DHX8, ABclonal, A14724; anti-RBM22, ABclonal, A10025; anti-SNRNP200, ABclonal, A6063; anti-LMNB, ABclonal, A16685) to immunopurify the respective RBP complexes. The complexes were washed with RIP buffer, and the samples treated with proteinase K buffer (10% SDS and 10 mg/ ml proteinase K in RIPA buffer) for 30 min at 37°C with shaking. RNA was further isolated using phenol:chloroform:isoamyl alcohol (125:24:1) solution, precipitated using isopropanol, and re-suspended in RNase-free water.

### Library constructions, RNA sequencing, and bioinformatics analysis

GBM4 cells were transfected with miR-10b inhibitor, U6 ASO #1, and corresponding control oligonucleotides for 72 hrs. The libraries have been constructed and sequenced by Novogene Corporation. Briefly, a total amount of 500 ng RNA per sample was used as input material. Ribosomal RNA was removed by Epicentre Ribo-zero™ rRNA Removal Kit (Epicentre, USA), and the libraries have been generated using NEBNext® Ultra™ Directional RNA Library Prep Kit for Illumina® (NEB, USA). The libraries have been purified using the AMPure XP system, and their quality assessed by Agilent Bioanalyzer 2100. The libraries have been sequenced on an Illumina platform and at least 12G of paired-end reads per sample were generated (Novogene). The data filtering included removing adaptors, removing reads containing N > 10% (N represents base that could not be determined), the Qscore (Quality value) of over 50% bases of the read is <= 5. Alternative splicing analysis was performed by the software rMATS, a statistical method for robust and flexible detection of differential alternative splicing from replicate RNA-Seq data. It identifies alternative splicing events corresponding to all major types of alternative splicing from replicate RNA-Seq data. It identifies alternative splicing events FDR for differential splicing. These types include exon skipping (SE), alternative 5′ splice sites (A5SS), alternative 3′ splice sites (A3SS), mutually exclusive exons (MXE), and retained introns (RI).

### Luciferase reporter assay

Full-length human RNU6 sequence (110 nucleotides) was amplified by PCR and cloned into psiCHECK2 (Promega), downstream of Renilla luciferase, using XhoI and NotI restriction sites. Glioma cells were transfected with 100ng of the reporter constructs, including the original “empty” plasmid and U6 reporter, and 24 hours later transfected with either miR -10b mimic (final concentration of 25nM), miR-10b inhibitor (final concentration of 50nM), or the corresponding control oligonucleotides. Luciferase luminescence was measured using the Dual-Glo luciferase assay system (Promega, E2920) 24 hours after the last set of transfections.

### Statistical analysis

Details pertaining to all statistical analyses can be found in the figure legends.

## Acknowledgements

This work was supported by the R01 CA215072, R01 NS113929, and R21 AG060019 grants to AMK. We would like to thank Abdellatif El Khayari for assistance in analyzing CLEAR-CLIP data, Dr. Lien Nguyen for critical reading of the manuscript, and members of the Krichevsky lab for productive discussions. We thank the NeuroTechnology Studio at Brigham and Women’s Hospital for providing instrumentation access and consultation on data acquisition and data analysis.

## Author contributions

AMK and REL conceived and designed the study; REL and YZ performed experiments, data analysis, and visualization; ED and ZW contributed to data analysis; NMT contributed to CDC42 splicing analysis; MR, HS, RR, and EJU assisted with experiments; REL, YZ, and AMK wrote the manuscript. All authors revised and approved the manuscript.

## Data and materials availability

All data needed to evaluate the conclusions in the paper are present in the paper and/or the Supplementary Materials. Additional data related to this paper may be requested from the authors. The RNA-seq and CLEAR-CLIP data have been deposited to GEO and are accessible through GEO Series accession numbers GSE182280 and GSE182281.

## Expanded View Figure legends

**Figure EV1. Analysis of putative miR-10b binding sites in 5′ UTR, CDS, and 3′ UTR** of mRNA targets identified by CLEAR-CLIP, using a STarMirDB tool (Rennie, Kanoria et al., 2016), demonstrates the distribution of seed-based and seedless targets. Related to Figure 1.

**Figure EV2. Relative levels of miR-10b and U6 snRNA in glioma and non-glioma cell lines**. The qRT-PCR reactions were performed with equal RNA input and the Ct values are indicated.

**Figure EV3. MiR-10b is enriched in the spiceosome in glioma cells.**

A. Western blot analysis of LN229 and U251 nuclear fractions separated in glycerol gradient, demonstrates enrichment of the spliceosomal marker PRPF8 in the heavy fractions.

B. qRT-PCR analysis demonstrates miR-10b enrichment (Ct values < 25.5) in the corresponding heavy fractions in both cell lines.

C-F. SART3 and PRPF8 bind to AGO proteins in glioma cells. IP with either IgG (control), AGO2, SART3, or PRPF8 antibodies, followed by the Western blotting for the indicated proteins.

**Figure EV4. miR-10b regulates U6 snRNA but not U2 and U4 snRNAs in glioma cells.**

A. Representative denaturing Northern blotting of glioma cells and GSCs transfected with either U6 ASOs, miR-10b inhibitor or mimic, with probes specific for U2 and U4 snRNAs.

B. miR-10b mimic and inhibitor do not regulate 3′ UTR luciferase reporters bearing full-length U6 snRNA sequence in glioma cells. The data is presented as Renilla/Firefly relative luminescence and normalized to the corresponding values in cells not transfected with the oligonucleotides. n = 12; Graphical data are shown as mean ± SEM. ns, no significance.

C. qRT-PCR analysis of snRNA U6 levels in U251 cells transfected with either miR-10b inhibitor, mimic, or corresponding controls, and treated with Actinomycin D (mean ± SD, n=3). *P* values were calculated using two-way ANOVA. * *P* < 0.05.

D. U251 cells were treated with Actinomycin D followed by the qRT-PCR analysis of miR-10b and U6 levels (mean ± SD, n=3). *P* values were calculated using two-way ANOVA. **** *P* < 0.0001.

**Figure EV5. Effects of miR-10b and U6 snRNA on CDC42 alternative splicing and overall expression.**

A. Differential stability of CDC42 iso-1 and iso-2 variants. LN229 cells have been treated with Actinomycin D followed by qRT-PCR analysis of CDC42-iso1 and CDC42-iso2 variants.

B. qRT-PCR analysis of CDC42-iso1 or CDC42-iso2 levels in GBM8 cells transfected with either U6 ASOs, miR-10b inhibitor, mimic, or corresponding control oligonucleotides (mean ± SD, n=3). Fold-change in the expression of the isoforms is plotted relative to the corresponding control conditions. *P* values were calculated using two-tail unpaired t-test. * *P* < 0.05; ** *P* <0.01.

C. Western blotting analysis of the indicated proteins in GBM8 cells transfected with either U6 ASOs, miR-10b inhibitor, or mimic.

D. GBM8 cells were transfected with siCDC42-iso2, siCDC42-total, or control siRNAs, followed by the qRT-PCR analysis. * *P* < 0.05; ** *P* < 0.01; **** *P* < 0.0001; ns, no significance.

E. GBM8 cells were transfected with siCDC42-iso2, siCDC42-total, or control siRNAs, followed by Western blotting analysis.

F. The growth of GSC spheroids, untreated or transfected with CDC42-targeting siRNAs, have been analyzed. The number and size of GSC neurospheres have been calculated at day 7 after transfections (mean ± SD, n=3). *P* values were calculated using two-tail unpaired t-test. * *P* <0.5; ** *P* < 0.01; ns, no significance.

**Dataset EV1. MiR-10b chimeras identified by CLEAR-CLIP.** Related to STAR Methods. The table lists two most probable chimeric arms for each chimeric read along with their alignment and sequence information. Interacting chimeric regions were annotated with Transcript_ID, Gene_ID, Symbol, and RNA class. Exact genomic alignment positions are indicated for each chimeric arm. For identification of basepairing, IntaRNA software was used (Mann et al., 2017) and results are presented in the last two columns. More details are described in the Methods.

**Dataset EV2. Analysis of putative miR-10b binding sites** in mRNA (5′ UTR, CDS, and 3′ UTR) and ncRNA targets identified by CLEAR-CLIP, using the indicated prediction algorithms. P-predicted; V-validated.

**Dataset EV3. Changes in alternative splicing caused by miR-10b inhibitor.** GBM4 cells were transfected with miR-10b inhibitor and the corresponding control oligonucleotide for 72 hrs, and alternatively spliced mRNA isoforms profiled by deep RNAseq. Splicing alterations are summarized for five major categories (SE, MXE, A5SS, A3SS, RI). More details are described in the Methods.

**Dataset EV4. Changes in alternative splicing caused by U6 ASO 1.** GBM4 cells were transfected with U6 ASO1 and the corresponding control oligonucleotide for 72 hrs, and alternatively spliced mRNA isoforms profiled by deep RNAseq. Splicing alterations are summarized for five major categories (SE, MXE, A5SS, A3SS, RI). More details are described in the Methods.

